# GraMDTA: Multimodal Graph Neural Networks for Predicting Drug-Target Associations

**DOI:** 10.1101/2022.08.30.505168

**Authors:** Jaswanth K. Yella, Sudhir K. Ghandikota, Anil G. Jegga

## Abstract

Finding novel drug-target associations is vital for drug discovery. However, screening millions of small molecules for a select target protein is challenging. Several computational approaches have been developed in the past using Machine learning methods for computational drug-target association (DTA) prediction predominantly use structural data of drugs and proteins. Some of these approaches use knowledge graph networks and link prediction. To the best of our knowledge there have been no approaches that use both structural learning that offers molecular-based representations and knowledge graph-based learning which offers interaction-based representations for DTA discovery. Based on the premise that multimodal sources of information acting complimentarily could improve the robustness of DTA predictions, we developed GraMDTA, a multimodal graph neural network that learns both structural and knowledge graph representations utilizing multi-head attention to fuse the multimodal representations. We compare GraMDTA with other computational approaches for DTA prediction to demonstrate the power of multimodal fusion for discovery of DTA.

## Introduction

The drugs approved for various indications often have unintended activities or drug-induced adverse events (AEs). This is often attributed to a drug’s off-target effect, binding other known or unknown targets. While such off-target binding is one of the underlying causes for several drug-induced AEs, they can also be beneficial and may lead to drug repositioning or discovery of new indications for an approved drug. One of the important goals in drug discovery is to identify a drug lead that has minimal off-target effect with high binding affinity to selected target. Virtual screening is used for faster and economical discovery of potential drug-target associations (DTA), where computational screening of chemical libraries is performed using a target of interest.

In recent years, machine learning-based approaches are increasingly used for predicting DTA [1]. These methods typically compute drug and target similarities to predict the association or binding affinity scores. Recent advances in deep learning have enabled extraction of latent features directly from raw data. Previously to train deep learning models for predicting DTAs, the drug and target inputs, which are represented as SMILES^1^ and FASTA^2^ sequence respectively, are one-hot encoded. For instance, DeepDTA [2] and DeepConv-DTI [3] applied 1D convolution to the 1-hot encoded sequences and aggregate potential patterns from raw sequence information to predict drug-target affinity [2].

Text-based representations are fixed and are inflexible for additional atom or bond level features. Whereas graphs are flexible to incorporate such auxiliary information. Recently, graph neural networks have demonstrated their effectiveness for computational drug discovery [4]–[6]. Hence as an alternative, representing drugs as molecular graphs have been proposed for feature rich molecule representations. GraphDTA [7] and Graph-CPI [8], for example, have leveraged graph neural networks to learn the graph representation of molecules along with their additional atom and bond features [9], [10] for DTA predictions. Alternatively, DTA problem can also be considered as a link prediction task utilizing the bi-partite drug-target association network and their heterogeneous annotations. Through integration of various heterogeneous data, several methods have been proposed for drug-target relationship discovery [11]–[14]. NeoDTI, for instance, integrated heterogeneous network data from diverse data sources and utilized GraphSAGE [15] for predicting DTAs.

In majority of these approaches, drugs and target proteins are represented either as text or graph modality to learn their respective representations. In other words, the drug embeddings are learnt through text, molecular graphs or through its heterogeneous neighbors using graph neural networks. Similarly, protein embeddings are learnt through its FASTA-text sequence representation using convolutional nets or through its heterogeneous neighbors using graph neural networks. Most of these approaches however rely on single modal representation of drugs and proteins. Multi-modal representation learning on the other hand offers robustness when any of the modality is incomplete or corrupted and further the modalities act complementary to each other [16]. Hence, we report **Gra**ph Neural Networks for **M**ultimodal biomedical datasets to predict **D**rug-**T**arget **A**ssociation (GraMDTA).

We collect structural and knowledge graph associations of drugs and proteins from various sources. We harmonize the collected modalities of the data. Our approach involves a pretraining phase where we train graph neural network on knowledge graph. Next, we incorporate the knowledge graph-based embeddings in GraMDTA. Further, for drugs represented as molecular graphs, we use graph attention network. For proteins represented as FASTA sequence, we use convolutional neural network for learning representations. Therefore, unlike previous works [2], [3], [7], [13], [17], [18], GraMDTA learns representations for drugs and proteins from structures and their corresponding knowledge graphs. We aggregate the multiple modalities of drugs and proteins using multi-head attention weighting mechanism to learn relevant information while eliminating the noisy information cascades. To assess the effectiveness of our model, we use it for drug target association prediction and drug target affinity score prediction problems by postulating the model as classification and regression methods. We perform systematic analysis to test the robustness of our framework on benchmark datasets through ablation studies. Our performance comparison with baseline models and ablation studies suggests that, multimodal learning has significant advantage over single modal learning. Our results from case study on select druggable target proteins from the Illuminating the Druggable Genome (IDG) [19], [20] database highlights the translational utility of GraMDTA.

## Materials and Methods

In this section, we describe our collected datasets i.e., drug-target association dataset, chemical structure dataset, protein sequence dataset, and relevant heterogeneous networks. Then we explain the components involved in pretraining and GraMDTA.

### Datasets

Pretraining Datasets: Our pretraining datasets consists of knowledge graphs of drugs and proteins. We collected and normalized the heterogeneous associations of drugs and targets from publicly available datasets as described previously [11]. More specifically, we collected associations for all the heterogeneous entities from datasets such as DrugBank [21], RepoDB [22], DisGeNET (curated) [23], DrugCentral [24], STRING [25] and Pharos[19]. For entity normalization, we used DrugBank identifiers for drugs, UMLS concepts for diseases, UniProt identifiers for targets, and MESH identifiers for drug’s categories (Table 1).

**Table 1:**
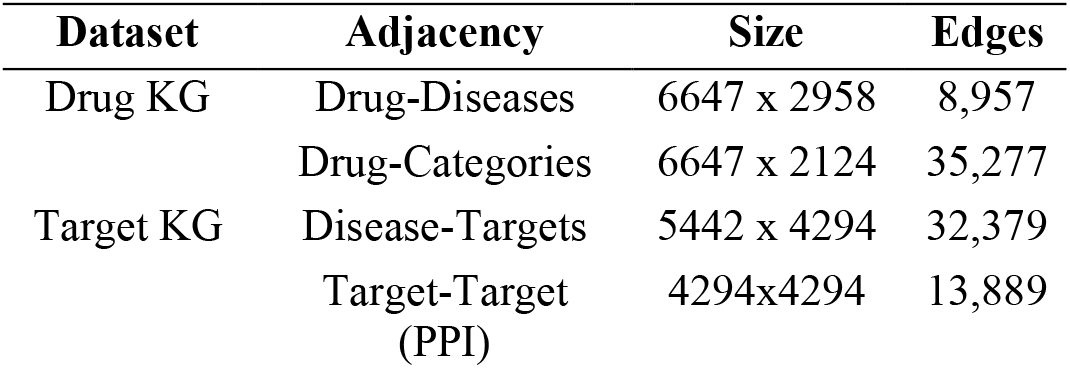
Knowledge graph information for drugs and target knowledge graph datasets.

| Dataset | Adjacency | Size | Edges |
| --- | --- | --- | --- |
| Drug KG | Drug-Diseases | 6647 x 2958 | 8,957 |
|  | Drug-Categories | 6647 x 2124 | 35,277 |
| Target KG | Disease-Targets | 5442 x 4294 | 32,379 |
|  | Target-Target (PPI) | 4294x4294 | 13,889 |

#### Benchmark Datasets

The benchmark datasets consist of known DTAs from 3 resources, namely, Drugbank [21], KIBA [26], and DAVIS [27]. The Drugbank dataset consists of drug-target binary associations. Out of 35,022 interactions in Drugbank, 17,511 interactions are positive and remaining are random negative interactions. KIBA and DAVIS datasets consists of affinity scores between drug-targets (Table 2). As part of pre-processing, we use RDKit software [28] and convert the SMILES representation of drugs to molecular graph representation where nodes represent atoms and edges represent bonds. The features which characterize the atoms and bonds are encoded in one-hot fashion similar to [29] and are used along with the molecular graph. For protein sequence, we one-hot encode the FASTA sequence text similar to previous works [2], [3], [7].

**Table 2:** Benchmark datasets and their statistics

| Dataset | #Drugs | #Targets | #Interactions |
| --- | --- | --- | --- |
| Drugbank | 6655 | 4294 | 35,022 |
| KIBA | 2068 | 225 | 116,350 |
| DAVIS | 68 | 379 | 25,772 |

### Pretraining

The pretraining stage consists of graph encoding and edge decoding phases. In the graph encoding phase, the feature information is aggregated from node neighborhood using graph neural network. In edge decoding phase, we perform link prediction based on the learnt node representations. In Figure 2, we show the encoding and decoding phases for drug and protein KG as generic representation.

#### Graph Encoding

In the encoding phase of pretraining, we use GraphSAGE for learning node representation. GraphSAGE aggregates the features of target node neighborhoods and concatenates the node’s current representation along with the aggregated neighborhood vector. This way the algorithm iteratively updates the node representation as:

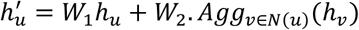

where *N*(*u*) is the neighborhood of node of u, *W*_1_ and *W*_2_ are the weights associated to learn representation, and Agg(.) is the aggregate function such as mean or sum. We use mean aggregator in our implementation.

Similarly, we use CNN to encode the drug or protein sequence i.e., molecule or FASTA sequence using 1D convolutions. We use adjacently with knowledge graph encoding to alleviate the representation issue for isolated nodes in knowledge graph. For drugs or proteins that lack interactions, structural information is used and representations are generated. Our convolutional encoding mechanism is similar to protein sequence encoding mechanism discussed in the GraMDTA section.

#### Edge Decoding

Here we define the edge decoding where we concatenate the representations of nodes and use feed forward layer to predict the link between them.

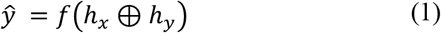

where ⊕ indicates concatenation of node embeddings *h*_*x*_ and *h*_*y*_, and *f*(.) is feed-forward neural network. We apply sigmoid activation to the output i.e., *σ*(*ŷ*), to predict the probability of link being associated between two nodes. We train the network using binary cross entropy loss function for both drug and protein knowledge graph datasets. The embeddings of drug and protein are extracted to further train with GraMDTA.

### GraMDTA

GraMDTA has three types of encoding networks with each encoder corresponding to a modality. The molecular graphs are represented as homogeneous networks; protein FASTA sequence is represented as text; and drug and target-related annotations as knowledge graph embeddings of heterogeneous networks (Figure 1). We define graph neural networks and convolutional neural net encoders according to each modality and then aggregate the encoded representations through a multi-headed attention mechanism. Further, we perform classification on the aggregated embeddings predicting DTA as probabilities or affinity score based on benchmark dataset. In the following sections, we define the encoders and corresponding loss function.

**Figure 1:**
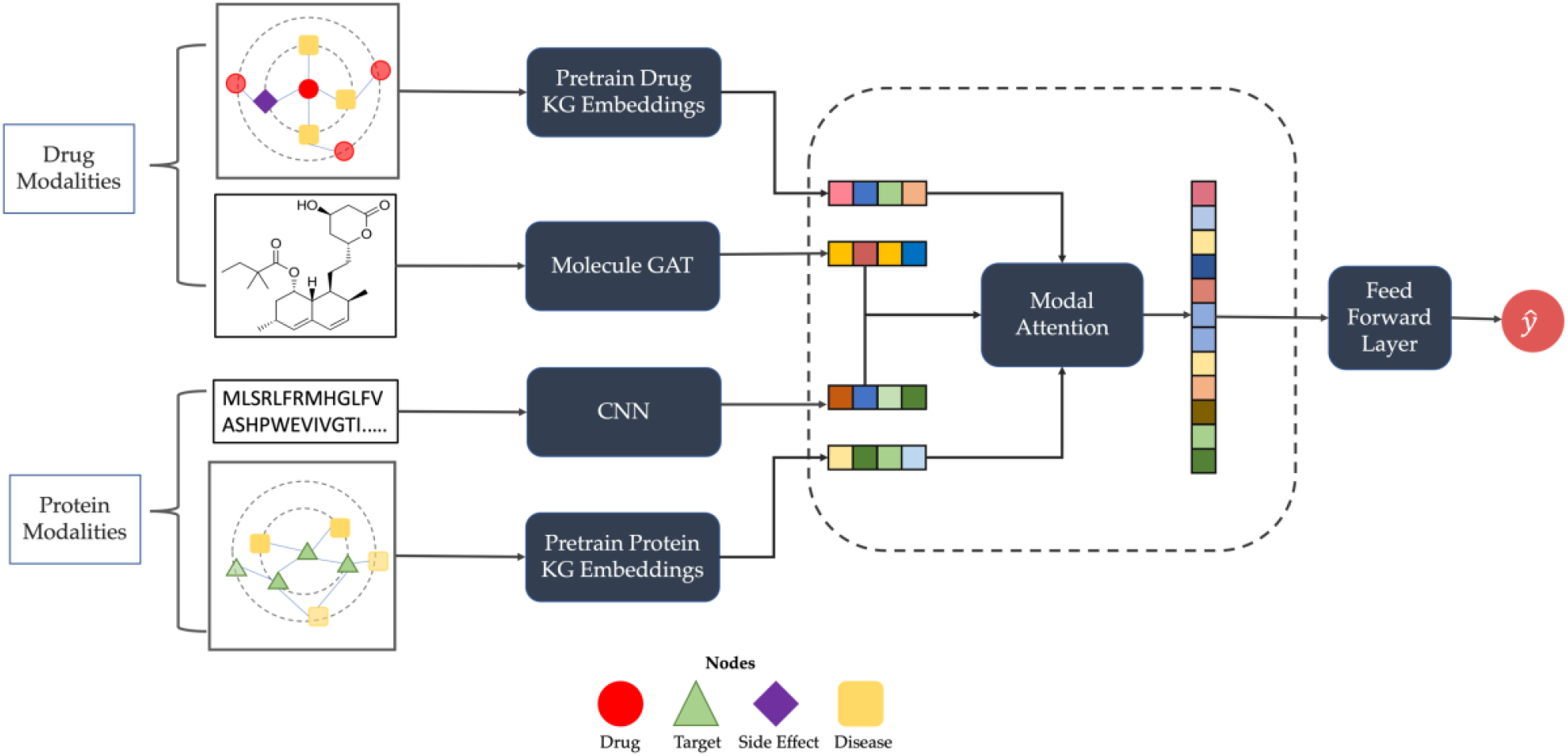
Overview architecture of GraMDTA for predicting association between drug and target protein using multiple modalities of the entities.

### Encoding Molecular Graphs

Molecular graphs are homogeneous networks where each node belongs to same node type (atoms) and edge type (bonds). To learn homogeneous graphs using neural networks, we define the GNN notations. A graph *G* = (*V, E*), represented as an adjacency matrix *A*, consists of *n* ∈ *V* nodes and *m* ∈ *E* edges. The sparse adjacency *A* can be preprocessed by converting to normalized Laplacian *Â* which is defined as a symmetric positive semidefinite matrix as 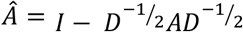. We use graph attention network (GAT) [30] to learn molecular representation which applies linear transformation for every node by a weight matrix *W*. Then for each node, attention coefficients are computed using their first-order neighbor nodes as follows

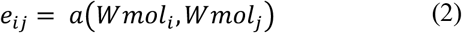

Where *i* is the target node index, *mol*_*i*_ is the target node (atom) embedding of graph *mol, mol*_*j*_ is the neighbor node (atom), *a* is a function to compute attention coefficients, and *e*_*ij*_ is node *j*’s influence over node *i* which quantifies the importance of the relationship. The scores are normalized using a SoftMax function as

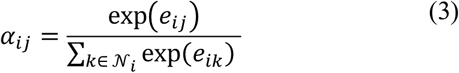

Using the normalized attention scores, node embeddings are computed using a non-linearity function *σ* as follows:

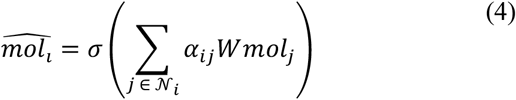

In this work, we use two layers of *GAT*(.) to generate graph embedding 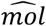. We further use a rectified linear unit (ReLU) as an activation function over the learnt embedding. The overall molecular graph embedding is simplified in the following notation

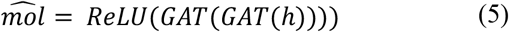

### Encoding FASTA Sequences

For 1D protein sequence, we one-hot encode and served as input for 1D-convolutions. The 1D convolutions are efficient in learning important local patterns within the sequence [2], [7]. The filters of convolutions slide through protein sequences effectively capturing relationships of an amino acids group. We denote the learnt representations of protein sequence (*p*) using convolution as

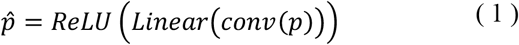

Where *conv*(.) is the convolutional operator, *Linear*(.) is the fully connected linear layer, and *ReLU*(.) is the activation function.

### Encoding Knowledge Graphs

The pretrained embeddings of knowledge graph are fed to feed forward neural network as:

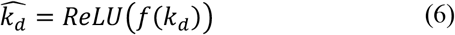

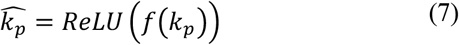

where *k*_*d*_ and *k*_*p*_ are knowledge graph-based representations of drugs and proteins, respectively.

### Modal Attention – Fusing Multiple Modalities

Here we discuss the modal attention to fuse multiple modalities. Before we dive into the details of fusing modalities, we first summarize the notations of each modality. For a drug, there are two modalities i.e., molecular graph and heterogeneous network associations. The molecular graph representation of the drug is denoted 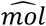 and the knowledge graph representation of the drug is represented as 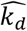. Similarly, the protein sequence has two modalities i.e., protein FASTA-text sequence and heterogeneous network associations. The protein sequence representation is denoted as 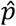 and the knowledge graph representation of protein is denoted 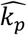.

We first concatenate the multiple modal representations of drugs and proteins. Then we use multi-head attention akin to attention in sentence transformers [31] where each modality acts as a word and the concatenation of the modalities is a sentence. With this analogy, we represent the modalities to multi-head attention as

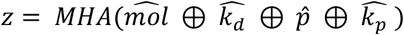

where ⊕ indicates concatenation and MHA(.) is multi-head attention which learns key features within and across modalities from different representation subspaces. Given each input as query *Q*, key *K*, and value *V*, the single head attention is defined as

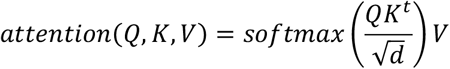

Since multi-head attention computes k attention heads, the final multi-head attention would be concatenation of all the attention heads (*α*_1_ … *α*_*k*_) with trainable parameter (*W*)i.e., *W*. (*α*_1_ ⊕ *α*_2_ ⊕ … ⊕ *α*_*k*_). We feed the final MHA(.) output i.e., 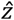 to 3-layer feed forward neural net along with dropout and ReLU activation function to predict the output.

### Loss Functions

Based on the task at hand, we define loss function for the benchmark datasets. For classification task using Drugbank dataset, we use cross entropy loss which is given as

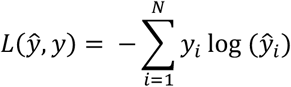

For regression task using KIBA and DAVIS datasets, we use mean squared error loss i.e.,

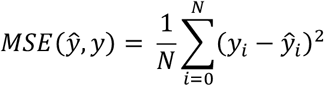

## Experimental setup and Results

In this section, we perform experiments on our collected benchmark datasets and compare with baseline models.

### Hyperparameter setting

Our GraMDTA model is implemented in PyTorch [32]. We use scikit to split the datasets into training, test, and validation sets. We consistently used the same sets to train baselines and our model for fair comparison. Additionally, we used Adam optimizer [33] where the learning rate is varied using cosine annealing with learning rate scheduler. We set the initial learning rate (η) set to 0.01. We trained the model for 1000 epochs and used early stopping when F1 score did not improve for 10 consecutive epochs. We trained our baselines similarly during comparison.

### Up-sampling Drugbank Dataset

While the associations between the drug and targets constitute the positive set, there is no negative set. To address the negative set issue, we pair random drug-target associations as negative set, similar to previous works [3], [7], [17]. This leads to a high-imbalance dataset depending on number of random parings being considered for each drug or target. Due to the high-imbalance of the associations, we limit to construct three datasets based on the up-sample ratios i.e., 1:1, 1:5, and 1:10. The 1:1 ratio has equal number of positives and negative associations where positive means there exists a drug-target association and negative association means no association between them. Then, we construct 1:5 where for every one positive association of a drug or target, we perform negative sampling of 5 associations with random pairing of target or drug respectively. We make sure that the randomly sampled drug-target association does not have any association across the training, validation, and test sets. Similarly, we construct a dataset having a ratio of 1:10 negative sampling where for one positive association we sample 10 negative samples.

#### Baselines

We compare our results with existing works with all the aforementioned experimental settings. Following are our baseline models:

##### DeepDTA [2]

A convolutional neural net model for learning SMILES and FASTA sequences which are one-hot encoded. In their methodology, they concatenate the representations of drug and target, then use fully connected layers to learn to predict the association.

##### HyperAttentionDTI [17]

This work learns representations using convolutions and applies attentions within and across drug and target sequences.

##### GraphDTA [7]

This work utilizes graph neural networks for molecules by representing SMILES as molecular graph and apply convolutions on FASTA sequence. Variants of GraphDTA are proposed using graph convolutions [34], graph attention [30], and graph isomorphism network [9] to aggregate the representations of molecular graph. In our experiments, we use both the variants to compare as baselines. Fully connected layers are further used to concatenate molecule and protein sequence embeddings to predict association.

#### Results

We evaluate our results on benchmark datasets i.e., Drugbank, KIBA and DAVIS. We segregate the results in to two task types i.e., classification and regression tasks.

To quantitatively evaluate the models, we adopt the metrics based on the task. For classification task, we use the standard metrics such as precision-recall (AUPR), area under receiver operating characteristic curve (AUROC) and F1 scores. To compute F1-score, we identify optimal threshold through based on elbow method of precision-recall curve. For regression task, we compute mean squared error (MSE), and concordance index (CI).

##### Task 1 – Classification on Drugbank dataset

We conduct experiments with our model using three up-sampling ratios as three experiment settings and compared results with the baseline models. The results in Table 3 show that, with increasing negative sampling size, the performance of the models declines. This suggests that performance of all models is affected with data imbalance. GraphDTA performed consistently better than baseline models with increase in negative sample ratio. Our model achieved superior AUPR and F1 scores across all the sampling ratios. Although, GraMDTA is marginally inferior with respect to AUROC in 1:1 and 1:5 sampling ratios, our model showed improvement over HyperAttentionDTI at 1:10 sampling. This demonstrates the model robustness with increase in negative samples.

**Table 3:** Classification performance comparison on Drugbank dataset

|  | 1:1 |  |  | 1:5 |  |  | 1:10 |  |  |
| --- | --- | --- | --- | --- | --- | --- | --- | --- | --- |
| Model | AUPR | AUROC | F1 | AUPR | AUROC | F1 | AUPR | AUROC | F1 |
| DeepDTA | 0.8439 | 0.8462 | 0.7815 | 0.6826 | 0.8759 | 0.641 | 0.5621 | 0.8666 | 0.5479 |
| GraphDTA - GCN | 0.773 | 0.8131 | 0.776 | 0.6497 | 0.8577 | 0.6172 | 0.5626 | 0.8687 | 0.5623 |
| GraphDTA - GAT | 0.8114 | 0.8312 | 0.7752 | 0.6773 | 0.8735 | 0.647 | 0.591 | 0.8837 | 0.62 |
| GraphDTA - GIN | 0.783 | 0.8101 | 0.7666 | 0.5757 | 0.8313 | 0.6566 | 0.5232 | 0.843 | 0.5428 |
| HyperAttention | 0.8766 | **0.8802** | 0.8099 | 0.7398 | **0.9079** | 0.6946 | 0.6706 | 0.9054 | 0.6386 |
| **GraMDTA (Ours)** | **0.8807** | 0.8767 | **0.803** | **0.7595** | 0.9041 | **0.7099** | **0.7245** | **0.9163** | **0.6861** |

##### Task 2 – Regression on KIBA and DAVIS datasets

Similar to Task 1, we conduct experiments evaluating the KIBA and DAVIS datasets as regression task. In Table 4, we show the performance results on both the datasets. With respect to KIBA dataset, GraMDTA achieved superior performance compared to all the baseline models. Our model achieved over 33% reduction in MSE and 6% improvement in CI when compared to the second-best reported work i.e., GraphDTA – GAT model. However, with respect to DAVIS dataset, HyperAttentionDTI had marginally improved performance with respect to MSE and CI. This suggests that there is room for improvement with respect to pretraining graph neural networks and GraMDTA encoding choices.

**Table 4:** Regression performance comparison on KIBA and DAVIS datasets (MSE: mean squared error; CI: concordance index)

|  | KIBA |  | DAVIS |  |
| --- | --- | --- | --- | --- |
| Model | MSE | CI | MSE | CI |
| DeepDTA | 0.149 | 0.852 | 0.151 | 0.919 |
| GraphDTA - GCN | 0.19 | 0.823 | 0.192 | 0.901 |
| GraphDTA - GAT | 0.103 | 0.88 | 0.166 | 0.915 |
| GraphDTA - GIN | 0.214 | 0.811 | 0.166 | 0.923 |
| HyperAttention | 0.231 | 0.81 | **0.108** | **0.947** |
| **GraMDTA** | **0.069** | **0.933** | 0.113 | 0.942 |

##### Ablation study

We conduct ablation studies on GraMDTA to understand the effectiveness and contributing components of the model by removing or substituting parts of the model. For the following ablation studies, we use 1:10 negative sampling from Drugbank dataset. The first row in Table 5, is the baseline score achieved from training Drugbank data with 1:10 negative sampling.

**Table 5:** Ablation results on GraMDTA

| Model | F1 score |
| --- | --- |
| GraMDTA | 0.686 |
| without pretrained-KG weights | 0.653 |
| with pretrained-KG weights only | 0.480 |
| Without Modal Attention | 0.66 |
| Pretraining GAT + GraMDTA | 0.665 |

First, we test our model by removing knowledge graph modalities of drug and target. We train the model with molecular graphs and FASTA sequence. This is equivalent to GraphDTA – GAT model along with modal attention component. The results in Table 5 suggest that removing knowledge graph embeddings (without KG) from the model led to 3% decrease in performance. Despite the removal of pretrained KG embeddings, the performance of GraMDTA without KG is still 3% better than GraphDTA – GAT, and 2% better than HyperAttentionDTI. This suggests that multi-head based modal attention have promising impact in learning representations.

Second, we remove the non-KG components i.e., molecular graph representation and sequence-based representation. While preserving modal attention component, we train the model using only pretrained KG embeddings. As shown in Table 5, without non-KG components, the performance of the model goes down by about 20%. This suggests that molecular graphs and protein sequence information are extremely vital for our GraMDTA model performance.

Next, we delete the modal attention component from GraMDTA and train the network. Although there is a marginal decline, our model performed better than the rest of the baseline models even after removing modal attention. Thus, demonstrating that attention improves the GraMDTA performance.

Finally, we change the pretraining encoder to Graph Attention Network (GAT) and pretrain the model for drug and protein knowledge graphs. We extract the embeddings and train GraMDTA with new embeddings. With GAT pretraining, there is 2% decrease in performance. Thereby, our results from ablation studies suggest that multimodal learning complements with structure-based learning i.e., SMILES and FASTA sequences, and further improves the performance of the model.

##### Interpreting Modal Attention

We extract modal attention for naltrexone (FDA-approved drug used to prevent relapses into alcohol or drug abuse) and its predictions. Our choice of the drug is arbitrary, and the goal of this analysis is to understand the attention contribution across multiple modalities. In Figure 3, ‘Molecule’ represents the molecular graph representation, ‘Drug KG’ represents drug KG representation, ‘FASTA seq’ represents sequence representation of protein, and ‘Protein KG’ represents protein KG representation. Each row in the subplot represents the attention scores computed through SoftMax normalization. The attention score suggests the importance of knowledge graph especially protein knowledge graph being consistently important for naltrexone and its protein associations. Similarly, molecular self-attention followed by FASTA sequence attention has shown to be consistently important. As seen in the figure, the attention scores for targets OPRD1, OPRM1, and OPRK1 have similar heatmaps suggesting identical binding associations. To validate this, we ran enrichment analysis on these genes using ToppGene Suite [35]. We found that the selected genes are enriched (p-value<0.01; FDR Benjamini and Hochberg) for gene ontology functions targeting opioid receptor activity (G-protein-coupled opioid receptor activity G-protein-coupled opioid receptor signaling pathway (GO:0038003) suggesting that GraMDTA can identify binding capacity for homologous structures and predict efficiently. This could be in part due to the high protein KG attention with each modality for the targets. On the other hand, although naltrexone and target SIGMAR1 have high molecule attention, the predictive probability is zero. This suggests that although attention signifies the importance of the input modalities, it alone may not be enough for predicting DTAs or potential mechanism of action [36], [37].

**Figure 2:**
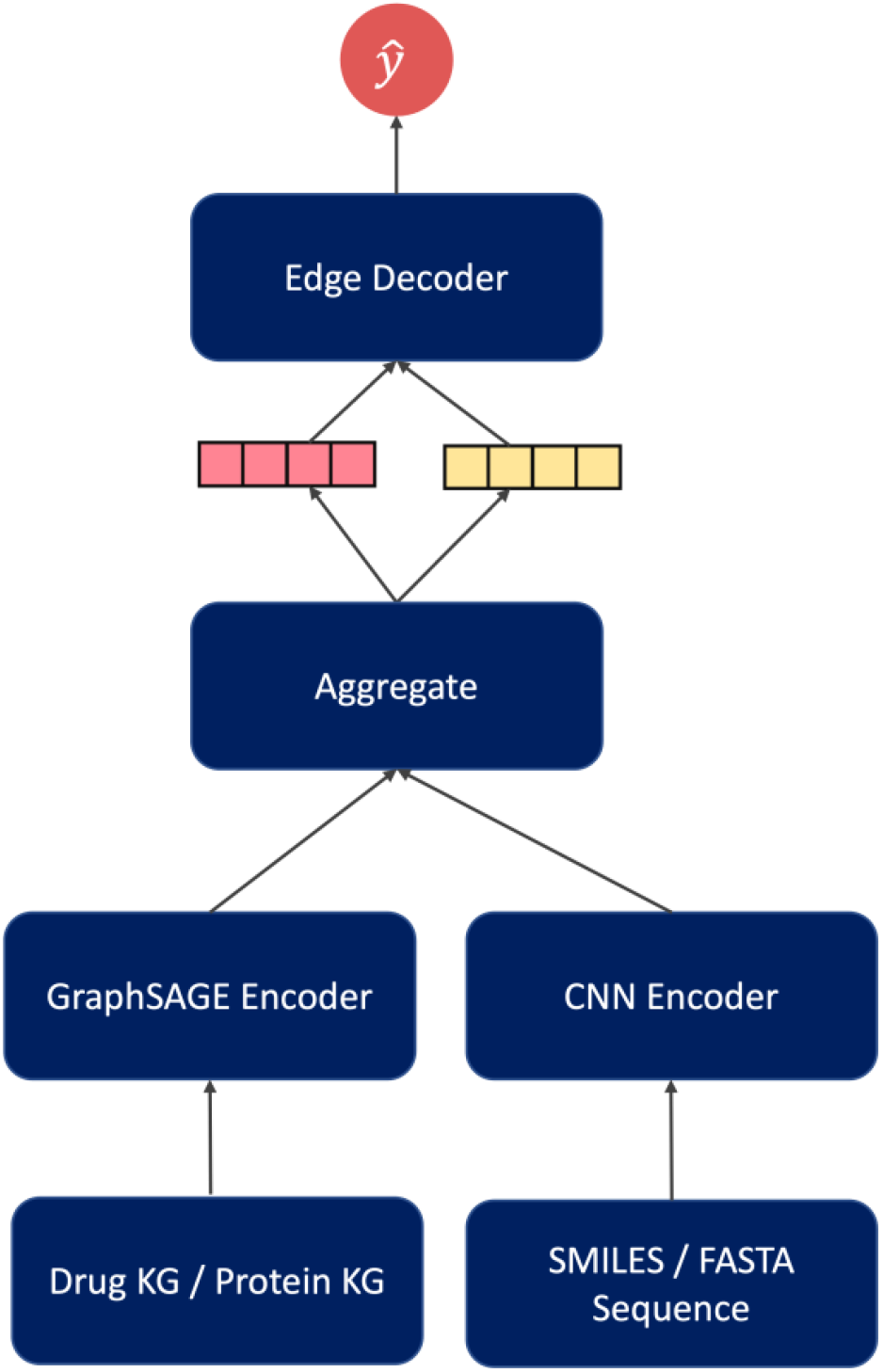
Overview architecture of pretraining model for training the knowledge graph

**Figure 3:**
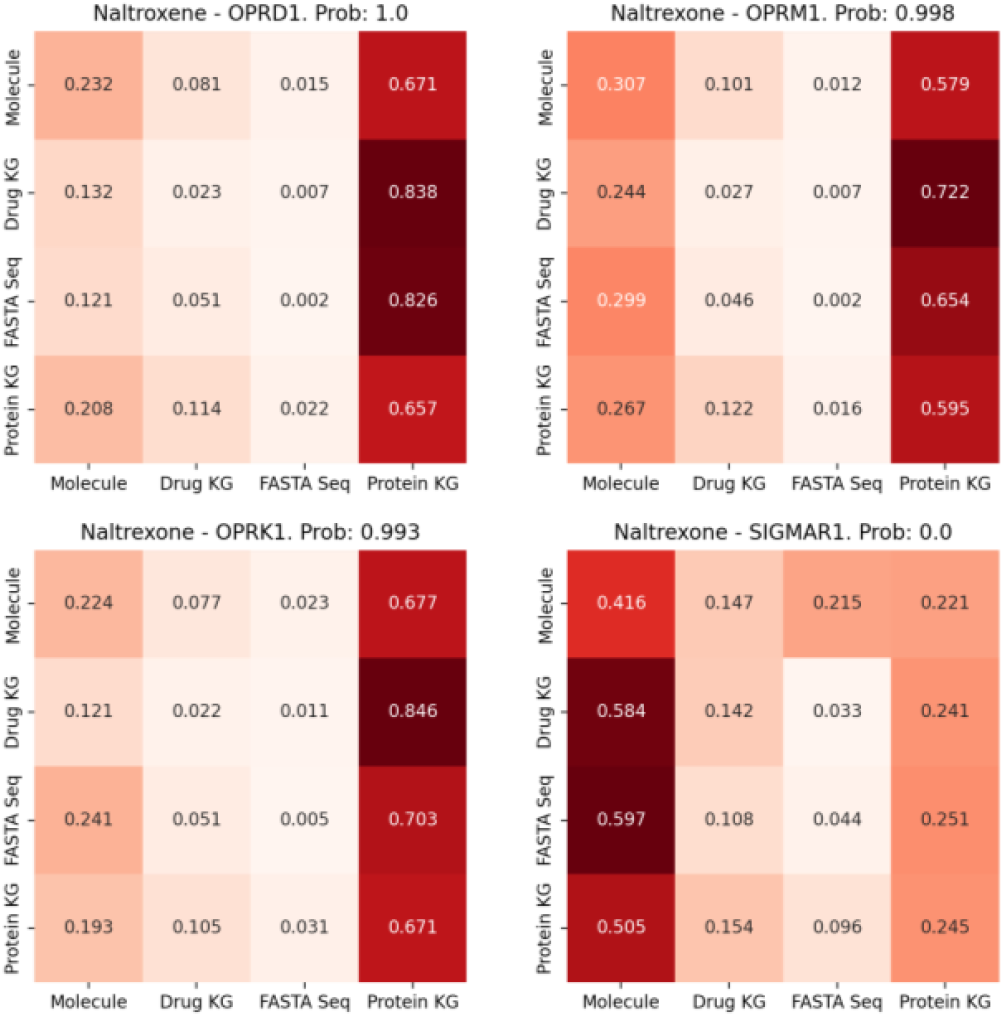
Attention scores for Naltrexone drug and its corresponding proteins ORPD1, OPRM1, OPRK1, and SIGMAR1. ‘Prob.’ in the subplot title indicates probability of drug and protein association.

## Case study

To demonstrate the translational utility of GraMDTA, we selected minoxidil drug and its target predictions as our case study. Minoxidil was initially developed to treat high blood pressure or hypertension [38]. Later, the drug was repositioned to treat hair loss or alopecia [39]. In Figure 4, we show the network visualization of minoxidil predictions consisting of targets which are both known and predictions. We performed enrichment analysis on targets predicted with high confidence (probability score >= 90%) using ToppGene suite [35]. Our results from ToppGene enrichment show that GraMDTA-predicted targets not only recover known indications but also potential candidates for repositioning. For example, enrichment analysis of GraMDTA-predicted minoxidil targets showed hypertension and alopecia, known and repositioned indications of minoxidil, respectively. Additionally, among the target functional enrichment are other diseases or biological processes such as arterial stiffness, breast cancer, and cognition suggesting the repositioning potential of minoxidil for other diseases. Indeed, literature review showed published studies in support of these. For instance, a recent study using breast cancer cells, reported that minoxidil, a known potassium channel opener, could inhibit cellular invasiveness in breast cancer. This study also reported that in combination with ranolazine, another ion channel blocker, minoxidil was synergistically effective as an anti-metastatic agent [40]. Another study on minoxidil revealed that the drug helps in reducing vascular or arterial stiffness [41], known to be associated with early-onset cognitive impairment and dementia [42], [43]. Likewise, modulation of potassium channels is known to play an important role in the regulation of memory processes [44]. Although, this example demonstrates the utility of GraMDTA for discovery of potential drug repositioning opportunities, it should be noted that mere enrichment of disease or phenotype does not always suggest a novel indication and may also suggest potential drug-induced adverse events. For instance, while multiple evidence suggest benefits of minoxidil in cognitive disorders, there are also conflicting reports in the literature reporting that minoxidil may cause cognitive impairment issues [45]. Nevertheless, the discovery of novel drug-phenotype associations through prediction of DTAs using GraMDTA and similar approaches are powerful in formulating translation hypotheses for drug repositioning or characterizing drug-induced adverse events.

**Figure 4:**
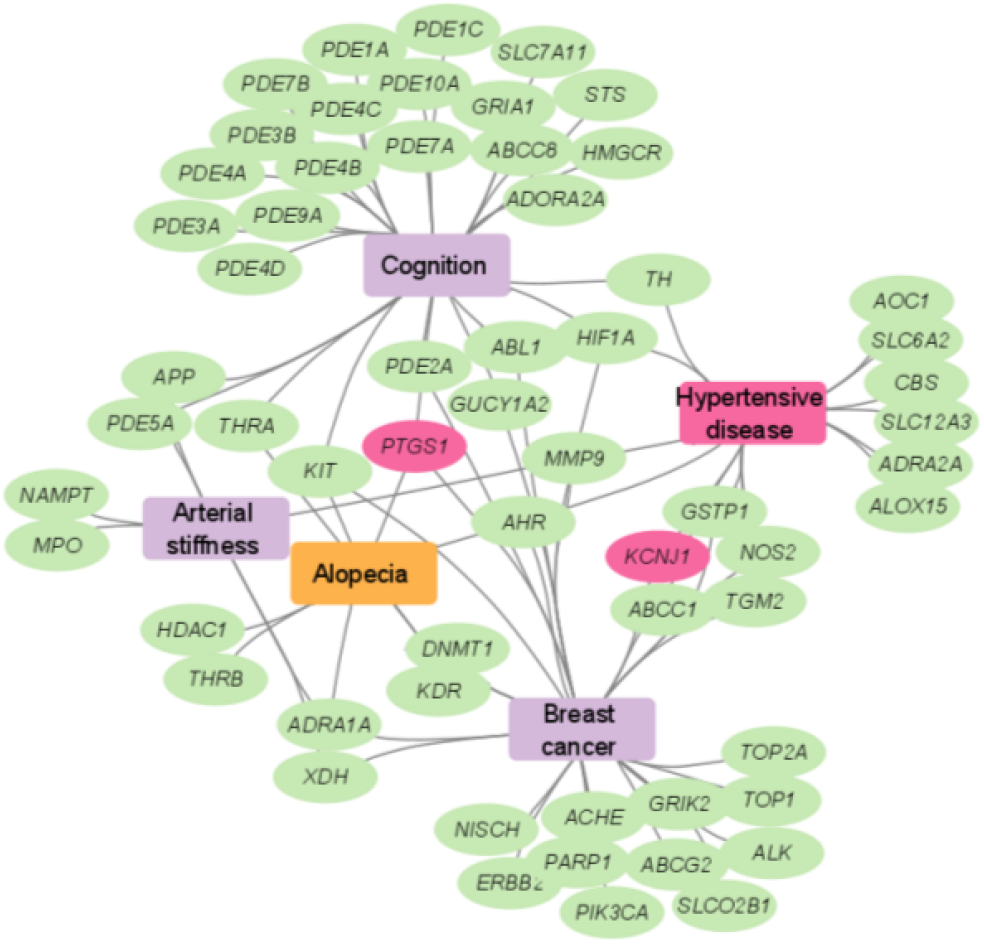
ToppGene enrichment analysis of targets predicted for minoxidil by GraMDTA. The rectangle nodes indicate indications (or diseases). The oval nodes indicate genes which are predicted with high probability (≥90% probability score). Pink-colored ovals are known targets of minoxidil. Approved known and repositioned indications (Hypertensive disease and alpoceia) are highlighted in pink and orange colors respectively. The other purple-colored rectangles represent potential novel indications (breast cancer, arterial stiffness, and cognitive disorders).

## Conclusion

In this paper, we report GraMDTA, a novel multimodal graph neural network based screening for discovery of drug-target associations. When compared using benchmark datasets, multi-modal aggregation outperformed other existing approaches. Our ablation studies further suggest that GraMDTA is robust in performance and demonstrate that supplementing pretrained knowledge graph representations to structural representations can improve the performance of the model. Our case study using minoxidil demonstrates that GraMDTA predictions could be useful for hypothesizing drug-phenotype associations (drug repositioning or drug-induced adverse events). Although pretraining knowledge graph using GraphSAGE supplemented GraMDTA, our pretraining could be further improved. An important future direction we plan to pursue is utilizing graph neural networks which are heterogeneous content aware [46]. More specifically utilizing context-specific gene expression (from human patients or animal models) information, and aggregating feature vectors accordingly. Secondly, we plan to expand our work on geometrical structures of protein as another modality in lieu with geometrical deep learning literature [47]–[49]. Finally, to encourage reproducibility of the work, we provide the source code and benchmark datasets at https://github.com/yellajaswanth/GraMDTA.

## Acknowledgment

This work was supported, in part, by the Cincinnati Children’s Hospital and Medical Center.

## Footnotes

1 Drugs are represented as SMILES (Simplified Molecular Input Line Entry System) which translates a 3-dimensional structure into a string of symbols.

2 Proteins are represented as FASTA sequences (amino acids) in a text-based format.

